# Cortical activity emerges in region-specific patterns during early brain development

**DOI:** 10.1101/2023.02.18.529078

**Authors:** Rodrigo Suárez, Tobias Bluett, Michael H. McCullough, Lilach Avitan, Dylan A. Black, Annalisa Paolino, Laura R. Fenlon, Geoffrey J. Goodhill, Linda J. Richards

**Affiliations:** The University of Queensland, Queensland Brain Institute; Brisbane, Australia; The University of Queensland, School of Biomedical Sciences; Brisbane, Australia; The University of Queensland, School of Mathematics and Physics; Brisbane, Australia

**Author notes:** Corresponding author, Linda J. Richards, AO, PhD FAA, FAHMS Washington University in St Louis School of Medicine, Department of Neuroscience, MSC 8108-12-8, 660 S. Euclid Avenue, St. Louis, MO 63110. These authors contributed equally to this work (Rodrigo Suárez, Tobias Bluett, Michael H. McCullough). Present addresses:* School of Computing and Eccles Institute of Neuroscience, John Curtin School of Medical Research, The Australian National University, Canberra, Australia (Michael H. McCullough). The Edmond and Lily Safra Center for Brain Sciences, The Hebrew University Jerusalem, Israel (Lilach Avitan). Washington University in St Louis School of Medicine, St Louis, USA #. **Author Contributions:** Conceptualization: LJR, GJG, RS Methodology: RS, TB, MM, LA, DAB, LRF, AP Investigation: RS, TB, MM, DAB, LRF, AP Visualization: RS, TB, MM, LA, GJG, LJR Funding acquisition: LJR, GJG, RS Project administration: LJR, RS Supervision: LJR, GJG, RS, LRF Writing – original draft: RS, TB, MM, GJG, LJR Writing – review & editing: RS, TB, MM, LA, DAB, LRF, AP, GJG, LJR. **Competing Interest Statement:** Authors declare that they have no competing interests. **Classification:** BIOLOGICAL SCIENCES, NEUROSCIENCE.

**Keywords:** cortical development, activity pattern, spontaneous activity, GCaMP6s, marsupial

## Abstract

The development of precise neural circuits in the brain requires spontaneous patterns of neural activity prior to functional maturation. In the rodent cerebral cortex patchwork and wave patterns of activity develop in somatosensory and visual regions, respectively, and are present at birth. However, whether such activity patterns occur in non-eutherian mammals, as well as when and how they arise during development remain open questions relevant to understand brain formation in health and disease. Since the onset of patterned cortical activity is challenging to study prenatally in eutherians, here we offer a new approach in a minimally invasive manner using marsupial dunnarts, whose cortex forms postnatally. We discovered similar patchwork and travelling waves in the dunnart somatosensory and visual cortices at stage 27 (equivalent to newborn mice), and examined progressively earlier stages of development to determine their onset and how they first emerge. We observed that these patterns of activity emerge in a region-specific and sequential manner, becoming evident as early as stage 24 in somatosensory and stage 25 in visual cortices (equivalent to embryonic day 16 and 17, respectively, in mice), as cortical layers establish and thalamic axons innervate the cortex. In addition to sculpting synaptic connections of existing circuits, evolutionarily conserved patterns of neural activity could therefore help regulate early events in cortical development.

**Significance Statement:** Region-specific patterns of neural activity are present at birth in rodents and are thought to refine synaptic connections during critical periods of cerebral cortex development. Marsupials are born much more immature than rodents, allowing the investigation of how these patterns arise in vivo. We discovered that cortical activity patterns are remarkably similar in marsupial dunnarts and rodents, and that they emerge very early, before cortical neurogenesis is complete. Moreover, they arise from the outset in different patterns specific to somatosensory and visual areas (i.e., patchworks and waves) indicating they may also play evolutionarily conserved roles in cortical regionalization during development.

## Main Text Introduction

Early neuronal activity plays an important role in driving the development of neuronal networks prior to functional maturation (1-7). The coordinated firing of neuronal ensembles during early stages of development has been detected using calcium imaging or electrophysiological recordings across a wide range of invertebrate and vertebrates species, including the fruit fly and zebrafish (8-10). Converging evidence suggests that early network activity in the mammalian brain regulates developmental processes including neurogenesis, neurotransmitter specification, cell migration, axon growth and synapse formation (11-18). However, precisely how these patterns affect early formation of the cerebral cortex remains largely unknown, as only mammals evolved a cerebral cortex and most of our knowledge about prenatal stages comes from in vitro studies. Outstanding questions include the extent to which early patterned activity is stereotypically region-specific, when and how they arise in development, and how conserved they are across mammalian species.

In newborn rodents, distinct patterns of spontaneous neural activity are present in the somatosensory and the visual cortices (SS and VIS, respectively) that are critical for the formation of topographic connections, and disappear by the end of the second week when the bulk of cortical connections are established. In VIS propagating patterns of activity are initiated by retinal waves and processed along visual areas, crucially priming for visual circuit function prior to eye opening (19-21). On the other hand, the SS of newborn rodents displays discrete patterns of non-propagated patchwork-like activity which, similar to waves in VIS, are driven by peripheral and thalamic spontaneous activity (22-24). Similar developmental features of cortical activity have been described in other mammals, including carnivores and primates, suggesting that shared mechanisms of functional and structural maturation might be at play (5, 6, 25-27). However, the fact that neural activity begins well before birth in eutherians has precluded detailed studies of its onset in vivo.

Live imaging of preterm mice delivered by C-section at embryonic day (E) 18.5 (i.e., mice are viable one day before term), has revealed that cortical activity relies on peripheral and thalamic input (1, 7). Cortical activity has also been recorded in vivo at earlier stages in acute preparations where mouse embryos remain attached to the maternal placenta (28, 29), or via a chronic implantation of an abdominal window in mothers (30). However, details of how activity patterns first emerge in the cortex remain largely unknown, as these approaches are highly invasive, usually involving maternal anesthesia that disrupts neural activity, and/or have low temporal and spatial resolution due to a combination of restricted choice of activity reporters, poor optical access, and little control over movement artifacts. To overcome these limitations, and to expand the phylogenetic repertoire of animal models of cortical development, here we studied a mouse-sized marsupial mammal, the fat-tailed dunnart *Sminthopsis crassicaudata*. Dunnarts are born extremely immature, at a developmental stage equivalent to mouse embryonic day 10 and human seventh gestational week, and undergo most of cortical neurogenesis and circuit development postnatally, within the pouch, making them easily accessible for mechanistic interrogations of brain development and evolution (31). We used in pouch electroporation of GCaMP6s, followed by live two-photon calcium imaging in the cortex of unanesthetized developing dunnarts, to examine the developmental dynamics of correlated patterns of activity.

We found that patchwork and wave patterns of spontaneous cortical activity emerge very early in somatosensory and visual areas, respectively, during stages of upper layer corticogenesis. These patterns are distinct from their onset, and mature with a rostral to caudal temporal delay as neuronal layers and thalamic input become specified, suggesting conserved mammalian features of cortical column formation prior to experience-dependent synaptic refinement.

## Results

We selectively expressed the genetically-encoded calcium indicator GCaMP6s (32) in pyramidal neurons via in-pouch electroporation (33), and performed in vivo two-photon microscopy in the cerebral cortex of unanesthetized dunnart joeys throughout stages of cortical formation (see Methods and *SI Appendix*). GCaMP6s was expressed using a piggyBac retrotransposase system (34) to transfect early radial glial progenitors and their progeny, consisting primarily of pyramidal glutamatergic neurons across all layers (i.e., not tangentially migrated cells such as interneurons; Figs. 1, S1). The few resident glial cells present at these stages were largely GCaMP6s-negative and there was no indication of increased cell death due to electroporation (Fig. 1a-d, and *SI Appendix* Figs. S1, S2 and Table S1), indicating that the signals were neuronal. We found that the translucent skull of developing dunnarts allows for minimally invasive monitoring of calcium activity across stages (S) S23-S27 (i.e., postnatal days (P) P27-P40), which are equivalent to mouse embryonic day (E) E15 to P4 (Fig. 1b, *SI Appendix* Fig. S1) (31). Two-photon images were captured between 50 – 200 μm below the skull/pial surface, with calcium signals therefore likely originating from both neuropil (including dendrites and axons) and cell bodies of pyramidal neurons.

**Figure 1:**
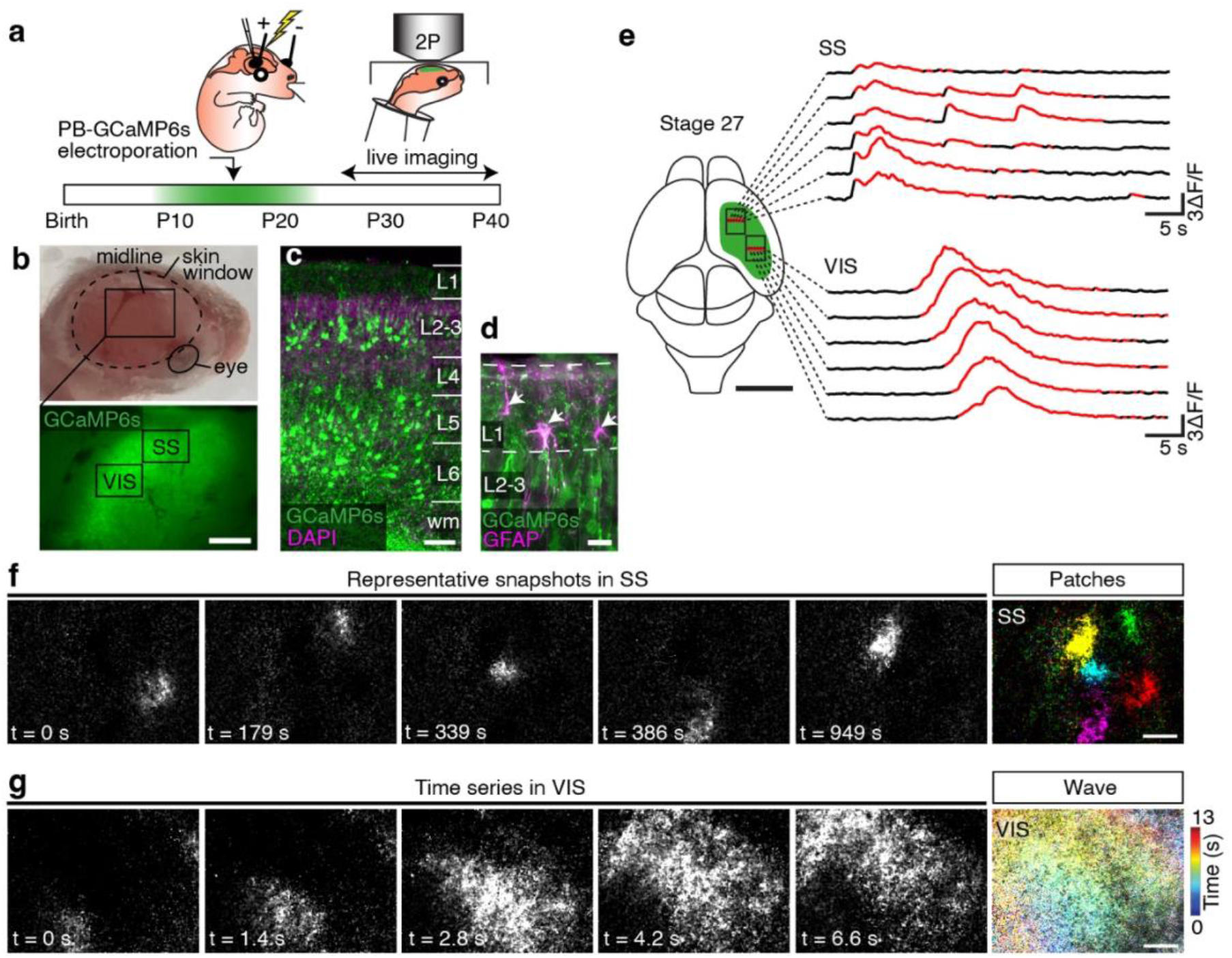
Conserved area-specific patterns of neural activity in the developing mammalian neocortex. **a**, Experimental design. Developing dunnart joeys were electroporated in-pouch with PB-GcaMP6s at P15, during the peak of pyramidal neuron generation, which extends from around P10 to P20 (stages 20 to 23, equivalent to mouse E12-15; grey area). Animals were then imaged using live two-photon (2P) microscopy between P21 and P40 (stages 23 to 27). **b**, Baseline fluorescence was readily apparent through joeys’ transparent skulls after opening a skin window. The square area is magnified below (inset) and indicates approximate regions of the somatosensory (SS) and visual (VIS) cortices imaged in this study. **c**, Coronal section through SS showing layered distribution of GcaMP6s-expressing pyramidal neurons (green) across layers (L) and white matter (wm) of an animal collected after live imaging at stage 27. **d**, High-magnification section of a stage 27 somatosensory layers 1 and 2/3 stained against glial fibrillary acidic protein (GFAP, magenta). **e**, Representative calcium traces (ΔF/F, significant events in red) of spatially contiguous medio-lateral regions (red bar) of SS (top) and VIS (bottom) of a dunnart at stage 27. Similar to equivalent-stage perinatal mice, activity in the dunnart neocortex is highly area-dependent. Patchwork-type activity is evident in SS, and travelling waves in VIS. **f**, Representative snapshots of recordings in SS showing patchwork activity (color-coded on right). **g**, Continuous time series of a VIS recording showing a travelling wave (time-color projection map on the right). In **f** and **g**, rostral is to the right and lateral to the top. Scale bars: 500 μm in B; 100 μm in **c**; 10 μm in **d**; 2 mm in **e**; 200 μm in **f**, **g**.

Previous investigations in postnatal rodents have revealed that cortical activity is organized in region-specific patterns during early development (19-23). To determine whether patterned cortical activity also occurs in non-eutherian mammals, we first recorded activity in the developing somatosensory and visual cortices (SS and VIS, respectively) of S27 dunnarts, equivalent to early postnatal mice (31). We analyzed activity within a 16 x 12 grid (870 x 650 μm) consisting of 54 x 54 μm regions-of-interest (see Methods and *SI Appendix*) and found that the dunnart neocortex displayed region-specific activity patterns that were remarkably similar to the patterns reported in rodents (Fig. 1e-g). In the somatosensory cortex we found patchwork activity: non-propagating events of synchronized local ensembles of around 100-200 µm in diameter (Fig. 1e-f, *SI Appendix* Movie S1). In contrast, visual cortex activity was characterized by larger-scale travelling waves (Fig. 1e, g, *SI Appendix* Movie S2). We found that patchwork activity in SS and wave activity VIS was consistently region-specific in the dunnart cortex, and their respective spatiotemporal dynamics remarkably resemble the patterns of spontaneous activity present during the first postnatal week of mice in homologous cortical areas. We then set out to investigate when and how these patterns of activity first arise by examining earlier stages.

Cortical neurogenesis occurs postnatally in marsupials, and pouch-young joeys do not need the placenta to breathe, therefore dunnarts are ideally suited to examine cortical activity from stages equivalent to embryonic rodents in vivo. These experiments revealed that correlated calcium events are typically very rare at S23, but patterned activity becomes evident in the SS by S24 (equivalent to E15 and E16 mouse, respectively). These earliest activity patterns were characterized by infrequent and sparse events that nonetheless exhibited some modular structure expected of patchwork-type activity (Fig 2a). Patchwork activity was significantly better defined by S25 (equivalent to E17 mouse), and displayed a spatial organization that was dominated by such modular ensembles (Fig 2a). Consistently, average pairwise Pearson correlation between regions of interest (ROIs) significantly increased from S23 to S25 when ROIs were <200 μm apart; a distance roughly corresponding to the diameter of each “patch”, suggesting formation of functionally relevant spatial modules (Fig. 2b). We next quantified the structure of patchwork activity by estimating functional network connectivity based on partial correlations between ROIs using Gaussian graphical models (35-37). The size of the largest connected component in the resulting networks increased from a few ROIs at stage 23 to encompassing almost all ROIs by stage 25 (Fig. 2a, c). Mean node strength showed an overall increase in correlated activity over this period (Fig. 2c). We then applied Louvain community detection to the networksClick or tap here to enter text. to infer the presence and structure of neural ensembles (38), defined as clusters of correlated ROIs in the network model based on modularity maximization. We found that neural ensembles emerged and increased in number and size over development, while network structures became increasingly modular (Fig. 2c). A decrease in the mean shortest path length normalized by network size (i.e., relative shortest path length, Fig. 2c) suggests that the network structure in the dunnart SS becomes a more efficient substrate for information transfer over development. This progression is similar to patchwork activity in the mouse SS, which becomes increasingly synchronized during the first postnatal week, including larger non-propagating events by postnatal day (P) 9, before becoming sparser and desynchronized by P11-13 (22, 23). Our findings in the dunnart SS indicate that although patterned activity becomes evident by S24, it becomes dominated by functionally modular neural ensembles that resemble newborn mouse patchwork activity by stage 25 (equivalent to prenatal mouse E17).

**Figure 2:**
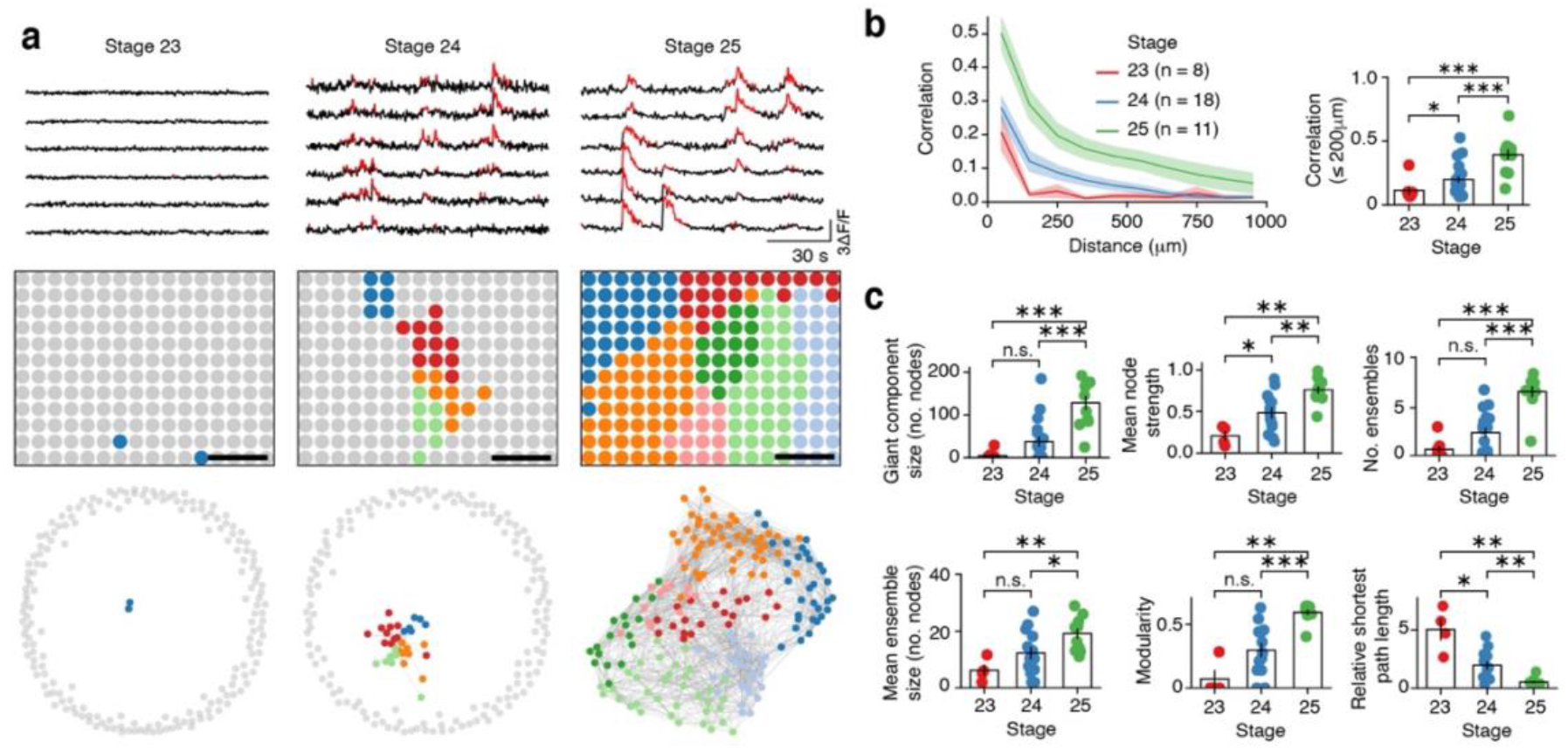
The onset and maturation of patchwork activity in the dunnart somatosensory cortex occurs between stages 23 and 25. **a**, Representative ΔF/F traces of contiguous regions of interest (ROIs, top rows), network nodes mapped to their spatial arrangement within the SS imaged region (middle rows, rostral is to the right and lateral to the top; scale bar, 200 µm), and functional connectivity networks estimated by Gaussian graphical modelling (bottom rows, color-coded nodes represent ROIs and edges represent non-zero partial correlations between pairs of ROIs), from three individual animals at stages 23, 24 and 25, respectively. Colors indicate distinct neural ensembles estimated by Louvain community detection. Grey nodes have zero partial correlation with all other ROIs. A single connected component (giant component) emerges during development, which encompasses the imaged region by stage 25. **b**, Pairwise Pearson correlation between ROIs as a function of distance increases over development (left) which is significant when averaged over distances less than 200 µm (right). Curves show mean correlation with respect to distance as first averaged for each animal then over all animals, shaded regions are the SEM, and individual data points in the bar plots are values for each animal. **c**, Properties of functional connectivity networks change over development, reflecting the emergence of patchwork-type activity. * p < 0.05, ** p < 0.01, *** p < 0.001.

Neural activity in the developing visual cortex (VIS) was characterized by wave-like patterns that displayed a diverse array of projection trajectories (Fig. 3, *SI Appendix* Fig. S3 and Movie S3). Similarly, waves are present along the visual pathway of eutherian newborns, including spontaneous waves in the retina, superior colliculus, thalamus and visual cortex, which are critical for functional maturation prior to eye opening and vision (2, 3, 19-21, 27, 39-42). However, when and how these waves begin in the cortex remains unclear. In dunnarts, we found that activity in VIS was first evident by S25 as infrequent and slow wave-like events, which then acquired clear wave features by S26, including significant increases in duration, propagated distance, size and area (Fig. 3b-c). Whilst waves seldom showed linear trajectories, they tended to be initiated rostro-medially and to propagate caudo-laterally at stage 26 (Figs. 3b, d; *SI Appendix* Movies S2 and S3). Waves had significant directional bias at stage 26 but not at stage 27, as they subsequently travel further and faster with more complex trajectories (Fig 3b, d and *SI Appendix* Movie S3). Velocity vector fields estimated by averaging wave trajectories from multiple animals (Fig. 3b, bottom rows) show complex patterns at stage 27 suggestive of anti-clockwise rotation over the imaged region similar to complex travelling waves observed in eutherians (43, 44). Importantly, these findings, as well as those for SS patchwork activity, remained consistent when using different sized ROIs for analysis, indicating that they are not sampling artifacts (*SI Appendix*, Fig S4). Taken together, these results indicate that the onset of patterned cortical activity in VIS occurs by S25, one stage after patchwork activity is evident in SS, and matures into complex travelling waves that parallel perinatal mouse VIS activity by S26.

**Figure 3:**
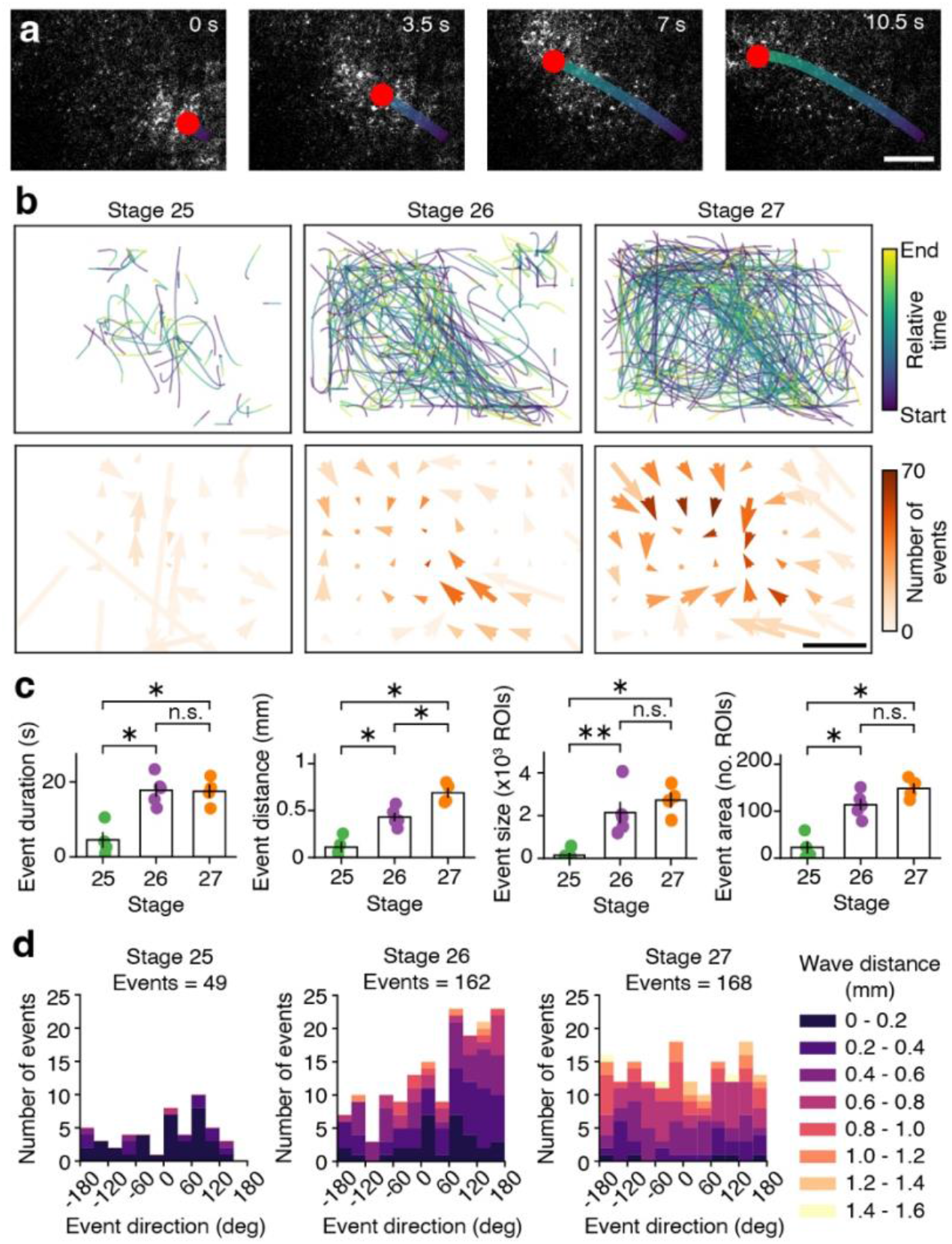
Onset and maturation of travelling waves in the dunnart visual cortex occurs between stages 25 and 27. **a**, Example of automated detection and tracking of travelling wave trajectories in VIS by connected components analysis and Kalman smoothing (see Methods and Fig S2). **b**, Top row: Trajectories for individual waves in VIS at stages 25, 26 and 27, pooled over all animals at each stage and color-coded by relative time. Waves with characteristic trajectory patterns emerge at stage 26 and become more complex by stage 27. Bottom row: Wave velocity vector fields over the VIS imaged region across development, corresponding to the wave trajectories above. The size and direction of each arrow corresponds to the average speed and direction, respectively, of waves travelling through each element of an 8 x 6 grid over the imaged region. Arrow color represents the number of waves that passed through each element of the grid. Directional structure appears at stage 26. Patterns at stage 27 show anticlockwise rotation over large sections of VIS. **c**, Event duration, distance, size (total number of ROI activations in the event) and area (number of ROIs spanned by the event) increased significantly from stage 25 to stage 26. Each data point is the average value over all events for an individual animal. **d**, Stacked histograms of event direction and distance. Direction is defined as the average along an event trajectory. At stage 25, detected wave events are few, only propagate over short distances (less than 200 µm, dark purple) and have no significant dominant direction. At stage 26, waves travel further and with statistically significant directional bias towards the upper-left quadrant of the imaged region (e.g., from rostro-medial to caudo-lateral). (Rayleigh test: p < 0.001; Rayleigh bimodal test: p = 0.03; Omnibus test: p = 0.001; Rao spacing test: p = 0.05; Hermans-Rasson test: p < 0.001). Wave distance continues to increase by stage 27 and there is no longer a single clear average direction of travel (no significant deviation from a uniform circular distribution). In **a**-**b**, rostral is to the right and lateral to the top. Scale bars: 200 µm in **a**, **b**.

Our findings indicate a delayed developmental maturation of cortical activity in VIS compared to SS, with the onset of SS activity occurring from S23 to S24, while in VIS it arises from S24 to S25, starting from very rare and low amplitude events that we termed pre-onset (Fig. 4a-b, and *SI Appendix* Movies S4 and S5). We sought to evaluate whether such early activity was distinct between SS and VIS, or instead it differentiates into patchworks and waves from shared features, by comparing t-SNE dimensionality reduction analyses (45) and mean Δf/f distributions for these early stages (Fig. 4c and *SI Appendix* Fig. S5). Interestingly, these rare pre-onset events already showed different features between SS and VIS, as well as at the subsequent stages of activity onset, suggesting that distinct functional circuits might be in place. As expected, SS and VIS activity were significantly different at subsequent stages 26 and 27, as shown by linear discriminant analysis (LDA; Fig. 5a). We then compared activity features between SS and VIS from individual animals across stages to evaluate how activity differentiates between areas. At early stages (S24 and S25) all measures in SS exceeded those in VIS (i.e., the direction of the first principal component) and, to a lesser extent, the difference between measures of patchwork and wave activity, respectively. This reflects the emergence of patchwork activity in SS and the overall delayed onset of activity in VIS. However, at older stages (S26 and S27) the variation in activity across regions was dominated by properties of wave activity in VIS (Fig. 5b, c and *SI Appendix*, Fig. S6). Notably, patchwork and wave patterns become more clearly defined across these stages, indicating that they do not represent sequential stages of maturation (e.g., patchworks are not precursors of waves), but rather they correspond to bona fide stereotypical region-specific patterns. These findings demonstrate that the measures we employed are sufficient to reveal distinct developmental trajectories of neural activity in SS and VIS, as early as patterns of correlated activity emerge. Notably, the timing of these inter-regional differences in maturation of patterned activity is consistent with a rostro-caudal gradient of cortical development across mammals, including molecular, cellular and connectivity features (46-51).

**Figure 4:**
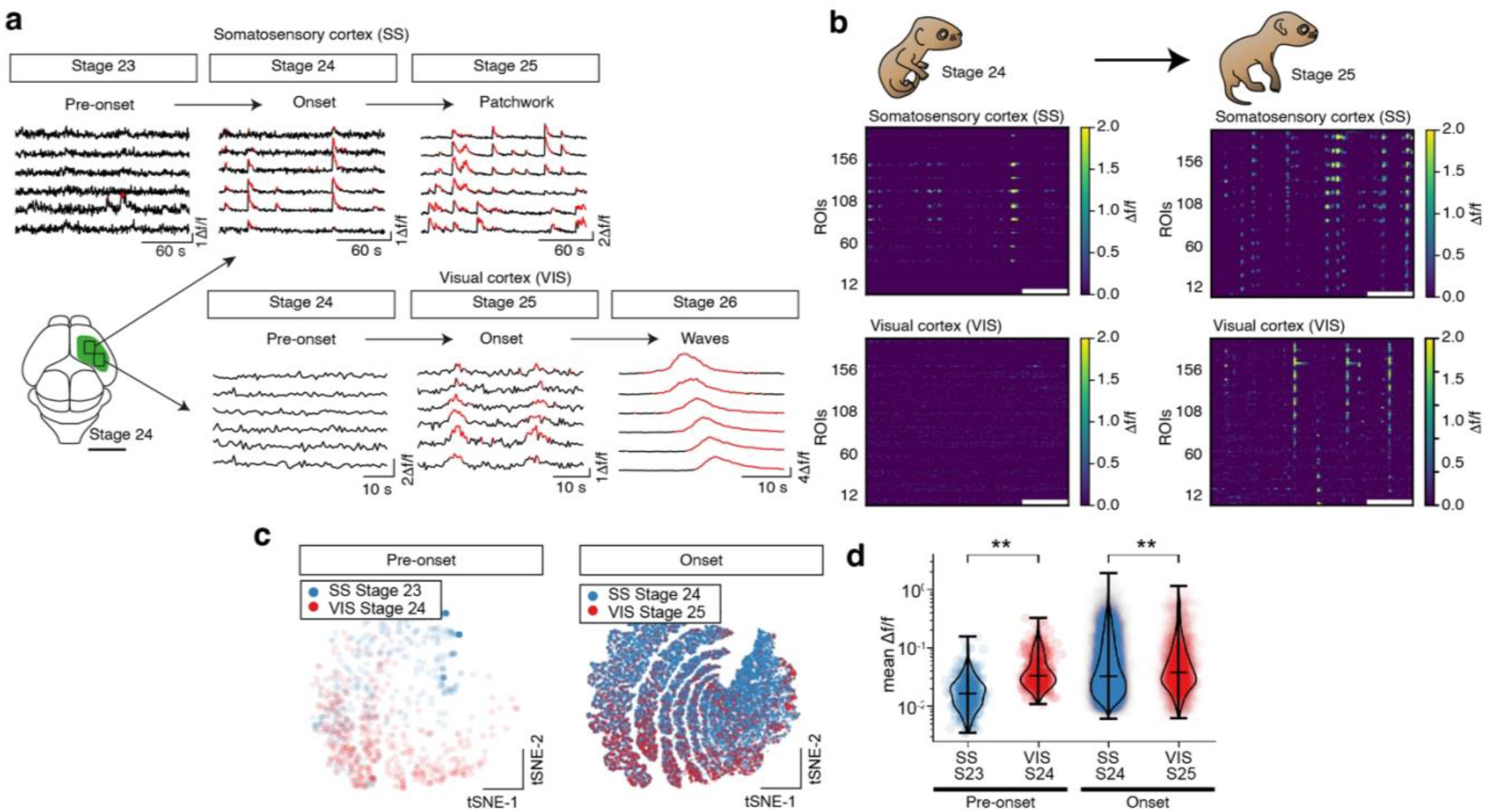
The emergence and specification of early neural activity in SS precedes that of VIS. **a**, Representative traces of activity within the somatosensory (SS, top) and visual (VIS, bottom) cortices across stages 23-26, including mostly silent pre-onset, activity onset, and clear patchwork or wave features becoming evident one stage later in VIS than SS. Schematics showing approximate imaging locations of a stage 24 brain. **b**, Representative raster plots of activity across ROIs from recordings of SS and VIS in one individual at stage 24 (left) and another at stage 25 (right) showing the broader delay in activity onset and maturation. Raster plots are masked to show only significant deviations from baseline fluorescence. **c**, Low-dimensional representation of pre-onset and onset activity in both areas (t-SNE, t-distributed stochastic neighbor embedding; each point represents an individual calcium transient per ROI) reveal differences from their onset (see Supplementary Fig. S5). **d**, Violin plots of mean Δf/f for individual events comparing SS and VIS for pre-onset and onset activity (mean +/-SEM; Wilcoxon rank sum test: **p < 0.01).

**Figure 5:**
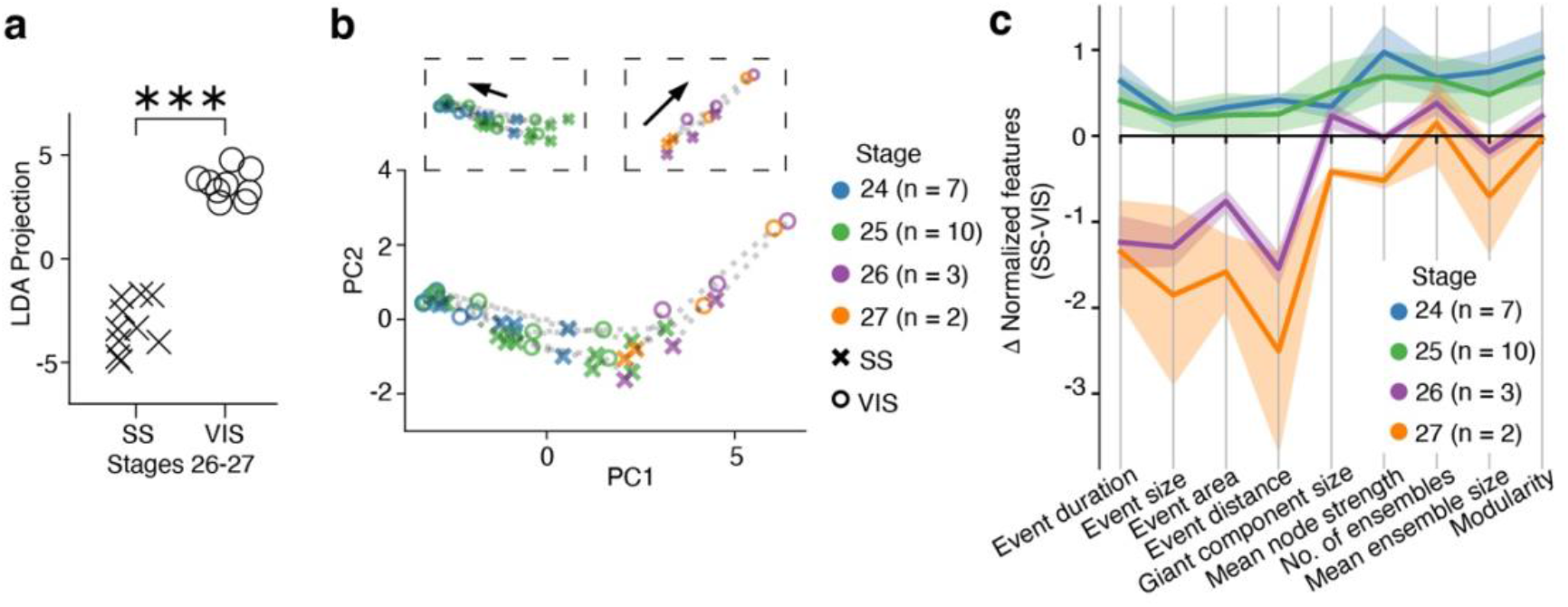
Developmental progression of correlated activity between somatosensory and visual cortex. **a**, Linear discriminant analysis (LDA) on a reduced feature set at late stages 26 and 27, when patchwork and waves have established, clearly separates data by region (MANOVA test: p < 0.001; each point represents one animal; see Methods and Supplementary Fig. S6). **b**, Principal component (PC) analysis of activity features from functional connectivity network analysis and wave event analysis for paired recordings of SS and VIS in single animals. Variables were scaled to zero mean and unit variance prior to computation of the PCs. Main axes show the projection onto the first two PCs which explain 93% of the variance in the data. Dashed grey lines between SS and VIS markers denote paired recordings from one animal. Inset panels show data separately for early stages (24 and 25) and later stages (26 and 27). Arrows show the average direction and magnitude of the difference between SS and VIS features for early and later stage recordings. The change in direction of the average feature difference over development suggests that patchwork and wave-like activity patterns in SS and VIS respectively do not develop in strict parallel. **c**, Feature difference profiles between activity in SS and VIS highlight distinctive properties of activity across developmental stages (mean +/-SEM). Activity features were computed by network analysis and wave analysis for paired recordings of SS and VIS in single animals and normalized for comparison. At early stages (24 and 25) all features have higher values in SS than VIS (i.e., positive values) with greater elevation for network features that characterize patchwork activity. At later stages (26 and 27) the difference between regions is dominated by the wave-like properties of activity in VIS.

Finally, to gain further insights on the neuroanatomical substrate that might underpin these differences of activity onset and maturation between areas, we compared cortical features across development in both areas. As expected, cortical width was consistently larger in SS than VIS across stages (Fig. 6a, b). Moreover, detailed layer characterization by immunolabelling of the intratelencephalic neuron marker Satb2 and the corticofugal neuron marker Ctip2, revealed that SS and VIS differ in their maturational dynamics, as quantified with a maturation index of the log ratio of upper layers (L2-4) over the proliferative compartment (ventricular zone) which undergoes a developmental shift between stages. This shift is consistent with maturational differences between areas (Fig. 6c), and might help explain the delay in onset of structured activity in VIS in relation to SS. Furthermore, detailed quantification of thalamocortical innervation across the cortex via immunolabelling of the vesicular glutamate transporter 2 (VGluT2), which is a specific marker of thalamic glutamatergic fibers (52-55), revealed further differences between areas, with SS initially showing higher values than VIS. As both areas differ in the absolute and relative size and proportion of cortical layers across stages, we normalized the columnar width, averaged the borders, and calculated VGluT2 intensity values at the middle 40% of each domain, minimizing potential confounds (Fig. 6d and *SI Appendix*). These differences were particularly salient at stages 24 and 25, when SS showed higher values than VIS initially in subplate and deeper layer compartments followed by higher innervation throughout layers, which then dissipated by S26. Note that, with exception of L1, no differences remain by S27, stage at which both areas display clearly structured patterns of activity (Fig. 1), and that a granular peak of thalamocortical innervation is not yet evident, as can be seen in layer 4 of adults (*SI Appendix* Fig. S7). The similar extents of VGluT2 expression at stage 27 between regions indicates that differences in neural activity patterns cannot be solely explained by differences in thalamocortical axon innervation at this stage. However, differences in thalamic input at early stages might correspond to differences in the onset of spontaneous activity, suggesting that not only intrinsic cortical maturation, but particularly as well thalamocortical input could explain the delay in activity onset in VIS as compared to SS.

**Figure 6:**
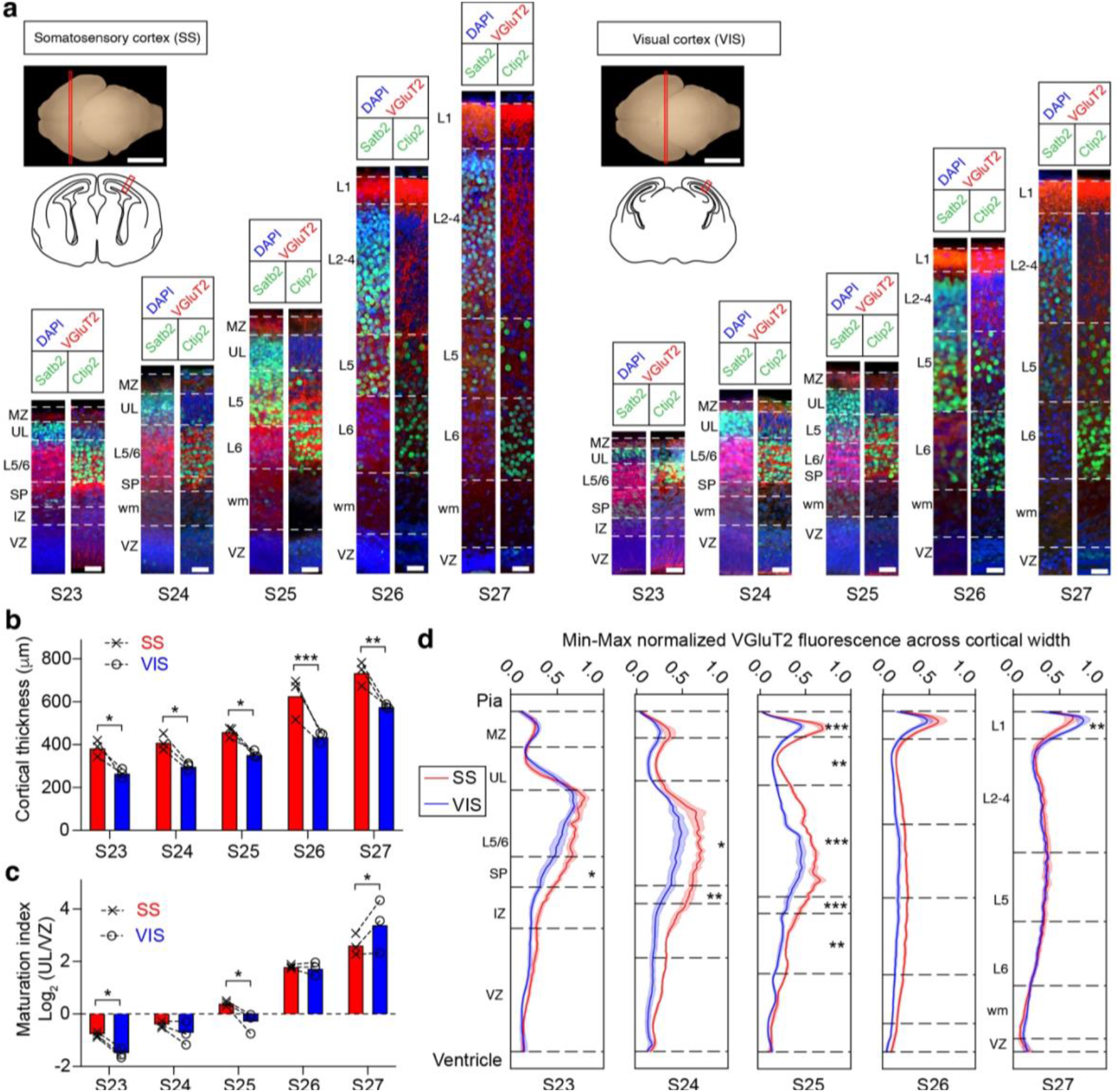
Differences in cytoarchitecture and thalamic innervation between somatosensory and visual cortices of dunnarts during stages 23-27. **a**, Representative coronal sections of somatosensory (SS, left) and visual (VIS, right) cortices stained with the thalamocortical axon marker VGluT2 (red), DAPI (blue), and either the upper-layer marker Satb2 or the deep-layer marker Ctip2 (green). Insets show approximate rostrocaudal extent (top) and a coronal schematic of the regions imaged in a red box (bottom) of a stage (S) 23 brain. Note the size and cytoarchitectural differences between SS and VIS across stages, taken from single animals per stage, and cortical thickness quantified as paired analyses in **b**, showing significantly larger columns in SS than VIS at all stages. **c,** Paired analysis of maturation index, quantified as a log_2_ ratio of upper layers (L) 2-4 and ventricular zone (VZ). **d,** Plot-profile quantifications of normalized VGluT2 fluorescence across layers (mean +/-SEM, n > 3 per condition) in SS (red) and VIS (blue). Dotted lines represent the extent of layers as depicted in **a**, averaged between SS and VIS for comparison, with asterisks representing significant differences between regions quantified within the middle 40% of each layer. **b-d**, Paired analyses using repeated measures two-way ANOVAs with Sidak’s multiple comparisons correction between areas per stage. * p < 0.05, ** p < 0.01, *** p < 0.001. IZ, intermediate zone; MZ, marginal zone; SP, subplate; UL, upper layers; wm, white matter. Scale bars in **a**: 2000 µm insets, and 25 µm in bottom sections.

## Discussion

The striking resemblance of region-specific patterned activity in dunnarts and rodents suggests that these network phenomena might perform similar functions in the establishment of cortical circuitry across mammals. In the rodent visual system, travelling waves of activity are driven by spontaneous activity in the retina, and support the formation of retinotopic circuits in the superior colliculus and primary visual cortex prior to vision (19-21, 40-42). Analogous travelling waves appear in the dunnart visual cortex at equivalent developmental stages and may therefore play comparable developmental functions. These travelling waves appear in the dunnart VIS 3-4 weeks before eye opening at stage 29 (31), are thus likely driven by spontaneous retinal waves, as in rodents. Likewise, in the rodent SS, patchwork activity has been implicated in the formation and fine-tuning of somatotopic barrels during a critical period of postnatal development, and distinguished from waves in VIS by potentially playing distinct roles shaping sensory pathways involved in discrete versus continuous perceptual maps, respectively (22-24). Although dunnarts lack morphologically conspicuous barrels in SS (but have a distinctive VGluT2-positive layer 4 evident in adults; *SI Appendix* Fig. S7), the remarkable similarity between dunnart and mouse patchwork-type activity suggests that it may also help establish discrete somatotopic maps in the dunnart cortex. In summary, our results demonstrate that equivalent region-dependent activity patterns are present in the marsupial visual and somatosensory cortices, and may therefore help establish functional circuits in these regions in a similar fashion to rodents. This hypothesis is supported by cytoarchitectural differences between SS and VIS, including absolute size, relative layering maturational states, and thalamocortical innervation, which broadly coincide with the emergence of structured neural activity in dunnarts, and by the fact that both waves and patchwork patterns activity are known to require intact thalamic and intracortical connections in early postnatal rodents (1-7, 19-24). The remarkable parallels in the spatiotemporal properties of cortical activity between dunnarts and mice could reflect independent origins in both lineages (i.e., homoplasy). However, additional similarities in cortical structure and function across therian mammals (46-48, 56, 57), instead favor the parsimonious alternative that these early patterns are an ancient feature of cortical development that has been conserved from common mammalian ancestors (i.e., homology). It is therefore likely that these cortical area-specific patterns of correlated activity play conserved and relevant roles in the formation and refinement of neural circuits in both species.

Our study also describes the ontogenesis of neural activity in the dunnart neocortex, which rapidly displays region-specific patterns at stages equivalent to prenatal eutherians. Notably, as similar patterns are already present at birth in rodents, our study represents a unique and minimally invasive system to further characterize the physiological mechanisms involved in the initiation and subsequent maturation of these neocortical activity patterns in vivo. Our findings of a protracted onset and maturation of activity in the visual cortex compared to the somatosensory cortex could be related to a rostro-caudal gradient of cortical development, as described across mammals (46-51). Alternatively, somato-motor functions may have an accelerated development in dunnarts, as they are critical to the survival of marsupial joeys from a very early age, with newborns needing to independently crawl and navigate into the pouch to locate a teat and begin feeding immediately after birth. Indeed, we recently reported that dunnarts have more mature neuronal markers than mice at the onset of corticogenesis (58), possibly also explaining our findings. Previous studies in rodents have similarly described a delayed developmental trajectory of network activity patterns in the visual cortex compared to the somatosensory cortex, with the visual cortex becoming responsive to light several days after the somatosensory cortex begins responding to tactile stimuli (21, 24, 59). It cannot be excluded that sparse cell assemblies could be active earlier than the stages we report here but are not undetected by our methods. Additional studies combining live electrophysiology and targeted manipulations might help provide additional details on the mechanisms of how activity develops. Similarly, comparative studies will be required to determine whether the differential timing in the onset of early activity patterns reported in the present study represents a phenomenon common to both marsupials and eutherian mammals, and whether perturbing such timing has any functional consequence.

Finally, our study establishes the fat-tailed dunnart as a model organism for the in vivo investigation of early neural activity patterns, which presents unique advantages when compared to rodents or other eutherians. In future studies, functional manipulations will likely reveal the circuit mechanisms that generate such region-dependent activity patterns, including the role of inputs from the periphery and subcortical structures such as the thalamus, as well as the developmental roles that such patterns may play in the precise formation and refinement of neocortical circuits.

## Methods

### Animals

All breeding and experimental procedures were approved by The University of Queensland Animal Ethics Committee and the Queensland Government Department of Environmental and Heritage Protection and were performed according to the current Australian Code for the Care and Use of Animals for Scientific Purposes (NHMRC, 8^th^ edition, 2013). Fat-tailed dunnarts were bred at the Hidden Vale Wildlife Centre, The University of Queensland as previously described (31). To detect the presence of joeys in the colony, the pouches of females were inspected every two-three days. The developmental stage of joeys was therefore P0-P3 days at inspection. The developmental stage was then confirmed according to head and body milestone features described previously (31).

### Plasmids

The calcium indicator GCaMP6s (Addgene 40753, MA) (32) was cloned into a piggyBac plasmid (kind gift of Joseph LoTurco, University of Connecticut, USA) (34). In this plasmid, GCaMP6s is flanked by piggyBac inverted terminal repeats. This pPbCag-GCaMP6s plasmid was co-transfected into developing dunnarts with a pCAG-PBase helper plasmid (kind gift of Joseph LoTurco, University of Connecticut, USA).

### In pouch electroporation

In pouch electroporation was performed as previously described (33). Briefly, when joeys were P9-P15, their mother was temporarily sedated by placement in a gas anesthesia induction chamber, where 5% isoflurane in oxygen was delivered at a flow rate of 200 mL/min. Sedated animals were transferred to a heating pad, where anesthesia was maintained via 2-5% isoflurane in oxygen delivered through a silicone mask (Zero Dead Space MINI Qube Anesthetic System, AAS, AZ). The joeys were exposed without removing them from the teat by carefully everting the pouch. 0.5-1 μL of a solution containing 1 μg/μL of pCAG-PBase plasmid and 1.5 μg/μl of pPbCag-GCaMP6s plasmid in 1X phosphate buffered saline (PBS; Lonza, Basel) and 0.0025% of the dye Fast Green (Sigma-Aldrich Co., MO; to visualize the location of injection) was injected into the lateral ventricle using a pulled glass pipette (WPI, FL) attached to a picospritzer (Parker Hannifin, NH). Five square pulses of 100 ms duration and 35 V were delivered at 1 Hz using 3 mm tweezer-like electrodes (Nepa Gene Co., Ichikawa) via an electroporation system (ECM830, BTX, Harvard Bioscience, MA), with the positive electrode placed above the developing cortex. In order to identify electroporated animals at later stages, animals were tattooed by scratching their paws with a hypodermic needle filled with green tattoo paste (Ketchum Mfg. Co., NY).

### Surgical procedure for in vivo imaging

Joeys were imaged between P20-P40. Mothers were anaesthetized and joeys were exposed as described above. Joeys were carefully removed from the teat by gently pulling, whilst squeezing the base of the teat with forceps. The joey was then restrained by wrapping its body in gauze and encasing it within a modified 1.5 μL centrifuge tube (Eppendorf, Hamburg), leaving the head exposed. Joeys were then anesthetized on ice for 4-6 minutes, or until unresponsive to stimuli. Topical local anesthesia (1% lignocaine; AstraZeneca, Cambridge) was applied to the skin on the scalp with a fine paintbrush, and the scalp covering the electroporated area was then removed. It was also critical to carefully remove the muscles superficial to the skull in order to prevent contractions that moved the sample during calcium imaging. Finally, a glass-bottomed petri dish (MatTek, MA) was fixed to the skull of the dunnart joey with cyanoacrylate glue (UHU, Bühl). The animal was then allowed to recover from ice anesthesia whilst the cyanoacrylate set for at least 30 mins.

### In vivo two-photon microscopy in unanesthetized dunnarts

To ensure consistent orientation of recordings, animals were aligned in the microscope such that the midline of the forebrain was parallel to the base of the field of view, with anterior to the right. Calcium imaging was performed using a Zeiss LSM 710 inverted two-photon microscope and Zen 2012 software (Carl Zeiss AG, Oberkochen). Samples were excited using a Spectra-Physics Mai TaiDeepSee TI:Sapphire laser (Spectra-Physics, CA) at an excitation wavelength of 940 nm. The emitted light was detected with a non descanned detector and a Plan Apochromat 10x/0.45 M27 objective (Carl Zeiss AG, Oberkochen). Time-lapse images were obtained from a cortical area of 870 x 650 μm for 10-30 mins at a sampling rate of 2.16 Hz and a pixel size of 2.18 μm.

### Tissue processing and histology

Immediately upon completion of two-photon microscopy, joeys were sacrificed by decapitation and their heads were immersed in 4% paraformaldehyde (PFA; ProSciTech, QLD) in PBS and post-fixed at 4 °C for 4-7 days. Heads were then transferred to 0.1% sodium azide (Sigma Aldrich, MO) in PBS. Brains were dissected and embedded in 3.3% Noble Agar (Thomas Scientific, NJ), and cut into 50 μm thick coronal sections using a vibratome (Leica Biosystems, Nussloch). Flat-mount sections were made by flattening the cerebral hemispheres after removal of subcortical structures, and sandwiching them between glass slides in 4% PFA prior to sectioning at 50 μm. To enhance green fluorescent protein (GFP) fluorescence, immunohistochemistry against GFP was performed on tissue sections from imaged dunnart brains. Sections were mounted on microscopy slides (Superfrost Plus, Thermo Fisher Scientific, MA), post-fixed in 4% PFA for 10 minutes, and washed 3 x 3 mins in PBS. Sections were then incubated in a blocking solution of 10% donkey serum (Jackson Immunoresearch Inc., PA) in PBS with 0.1% Triton-X (Sigma-Aldrich Co., MO) for 2 hrs. Sections were incubated with primary antibodies (polyclonal rabbit anti-GFP, 1:500, Abcam; rabbit anti-SATB2, 1:500, Abcam; rabbit anti-CTIP2, 1:500, Abcam; mouse anti-VGluT2, 1:500, Abcam; rabbit anti-cleaved caspase 3, 1:500, CST; chicken anti-GFAP, 1:1000, Abcam) in blocking solution overnight at room temperature. Sections were washed in PBS for 3 x 20 mins before being incubated with fluorescent secondary antibodies in 0.1% Triton-X for 2 hrs. Cell nuclei were stained with 4’,6-Diamidine-2’-phenylindole dihydrochloride (DAPI, 1:1000, Invitrogen, Thermo Fisher Scientific, MA) for 10 mins. Sections were cover slipped using ProLong Gold anti-fade reagent (Invitrogen, Thermo Fischer Scientific, MA) and stored in the dark at 4 °C until image acquisition. Wide-field fluorescence imaging was performed using a Zeiss upright Axio-Imager Z1 microscope fitted with an AxioCam HRm camera and a 5x objective with Zen 2012 software (Carl Zeiss AG,

Oberkochen). High resolution fluorescence images were obtained using a Diskovery spinning disk confocal microscope (Spectral Applied Research Inc, Ontario) built around a Nikon TiE body and equipped with two sCMOS cameras (Andor Zyla 4.1, 2048 x 2048 pixels) and controlled by Nikon NIS software (Nikon, Tokyo). All histology quantifications were done in FIJI (ImageJ, NIH), with Plot Profile function used to obtain grey values of the 16-bit images. The Multi-point function was used to demarcate layer boundaries based on Satb2, Ctip2, Dapi and VGluT2 channels in both hemispheres of SS and VIS of 3 or more animals per stage (S23-S27). A custom script in Python was used to linearly interpolate images to the size of largest sample per stage, and VGluT2 grey values were normalized between minimum and maximum values (min-max) into a 0-1 scale. Layer boundaries were averaged between areas within each stage, and the middle 40% of each interval was calculated for paired statistical tests. Imaging was performed in the Queensland Brain Institute and School of Biomedical Science’s Advanced Microscopy Facilities.

### Image processing

To investigate patterns of neural activity, the imaged region was partitioned into a 16×12 grid where each square region of interest (ROI) had a side length of 25 pixels (54 µm). We extracted the neural time series for each of the resulting 192 ROIs as the scaled and detrended change in fluorescence with respect to time:

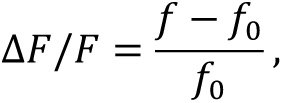

where 𝑓 is the mean fluorescence per frame of all pixels in an ROI, and 𝑓_0_ is a moving baseline for that same ROI. This baseline was estimated by applying a percentile filter to 𝑓 at the lower 10^th^ percentile with filter length of 100 seconds. We assumed that 𝑓_0_ would change slowly in time and therefore further smoothed our estimates of 𝑓_0_ using a moving mean filter with window size 100 seconds. If an ROI had a low baseline fluorescence (i.e., small 𝑓_0_) then it was treated as non-physiological data and the Δ𝐹/𝐹_0_ time series for that ROI was set to zero, effectively excluding these ROIs from subsequent analysis. Small 𝑓_0_ was defined as less than 10% of pixels having a non-zero fluorescence value over any 100 second window in the data.

#### Estimating binary neural activity

For each Δ𝐹/𝐹 we generated a binary time series which was 1 when significantly positive deflections in fluorescence occurred and 0 otherwise. This was computed per ROI by first using kernel density estimation to find the peak in the distribution of the Δ𝐹/𝐹 values. The bandwidth of the estimated distribution was computed using Silverman’s rule. We assumed that negative deflections from the peak represented a one-sided approximation of the noise in the data and estimated the standard deviation of the noise based on this. Deflections in the Δ𝐹/𝐹 time series were defined as significant if they were greater than 3 standard deviations above the peak of the estimated distribution.

#### Estimating functional connectivity networks

We investigated patterns of neural activity in each recording as functional connectivity networks where ROIs were treated as nodes. Network edges were weighted and undirected. Edge weights were computed based on partial correlations. Specifically, edge weight 𝑤_𝑖,𝑗_ was defined as:

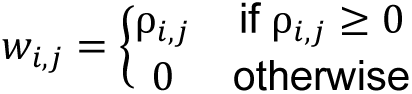

where 𝜌_𝑖,𝑗_ was the partial correlation between Δ𝐹/𝐹 time series between the between the 𝑖^th^ and 𝑗^th^ ROI. If 𝑤_𝑖,𝑗_ = 0 then nodes were disconnected. We only considered positive correlations because we sought to characterize coordinated ensembles of neurons that fired together, and neural spiking typically manifests as positive deflections in Δ𝐹/𝐹. Partial correlations were computed from the precision matrix Θ with elements θ_𝑖,𝑗_ as:

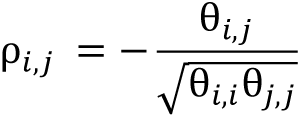

where Θ was estimated via Gaussian graphical modelling (37). Specifically, we used an implementation of the QUIC algorithm (60) in the Python package skggm (36) to solve the graphical LASSO formulation of the objective function for regularized maximum likelihood estimation of Θ (35). The regularization parameter λ was selected by the Bayesian information criterion (61) using a greedy hierarchical grid search initialized on 20 logarithmically spaced values of λ ∈ [0.01,1]. The grid was iteratively refined until the model error was less than 10^−3^ or 1000 iterations were reached. For ROIs with low baseline fluorescence which were set to zero during the computation of Δ𝐹/𝐹, we added low level Gaussian noise with standard deviation of 10^−6^ to avoid numerical instability in the algorithm.

#### Network visualizations and properties

Network properties were computed using the NetworkX package (62) for Python. See (63) for detailed definitions of the network properties summarized as follows. The giant component is the largest connected component in the network, that is, the largest set of nodes for which there exists a path between all pairs of nodes in that set. The mean node strength is equal to 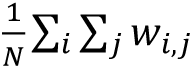 where 𝑁 is the number of nodes in the network, which in this study is 16 × 12 = 192 by the specification of the ROI grid. We defined the edge distance between nodes 𝑖 and 𝑗 as 𝑑_𝑖,𝑗_ = 1/𝑤_𝑖,𝑗_ for the computation of the mean shortest weighted path length. The mean node strength and relative mean shortest path length were computed on the giant component.

Estimation of neural ensembles was performed on the complete network using Louvain community detection (38) with the package Python-Louvain (https://github.com/taynaud/python-louvain). Louvain community detection estimates community structures (clusters of connected nodes) by finding an optimal partition of the network. The algorithm iteratively merges nodes together into a community if the merger provides a maximum increase in the network’s modularity (the strength of connections within modules compared to the strength between modules). This method is advantageous for comparing changes in community structure between networks (i.e., the number and size of communities) because it does not require the user to specify the number of clusters to search for, as is required by several other standard clustering algorithms such as 𝑘-means and spectral graph clustering. Instead, Louvain community detection iteratively merges nodes to form communities until there are no possible further mergers that will increase modularity, thereby automatically estimating the number of communities in the network. The iterative nature of the algorithm means that it can produce different solutions depending on the order by which nodes are considered for merging. Therefore, we ran the algorithm 100 times with different random node orderings for each recording, and ensemble properties including number of ensembles, ensemble size and modularity were averaged over the set of realizations.

Network visualizations were produced with the NetworkX package (62) using a force-directed graph layout (spring layout).

#### Tracking propagating spatio-temporal neural activity

We defined spatio-temporal events as contiguous volumes of binary neural activity in the temporal stack of imaging data from a recording. We considered the 16 × 12 × 𝑇 volume where the first two dimensions were the ROI grid and 𝑇 was the number of frames in the recording. We then populated each 1-dimensional column in the 𝑇 direction with the binary activity time series for the corresponding ROI. Events are connected components in the resulting binary volume. We used 6-connectivity for spatial adjacency in the 3-dimensional grid (i.e., elements can only connect along the primary axes; diagonal connectivity was not allowed).

We sought to exclude events which may have occurred by chance due to noise in spatially or temporally adjacent ROIs. Specifically, we kept only events which had volume, maximum cross-sectional area, and duration greater than a set of thresholds. Thresholds were computed for each recording separately by random circular shuffling the binary volume in the time dimension with a periodic boundary, where each ROI was temporally displaced by an independently drawn random offset. The distributions of volume, maximum cross-sectional area and duration were enumerated for 1000 shuffles and thresholds for significant events were set at the 95^th^ percentile of these null distributions.

To track the position of a significant event in the imaged region through its duration, we first created a binary mask of the event ROIs for the starting frame of the event. This mask was applied to the corresponding ROI grid of Δ𝐹/𝐹 values for this frame. The remaining non-zero ROIs were smoothed using a 2-dimensional Gaussian filter and the positions of the event for this frame was defined as the spatial peak of the smoothed Δ𝐹/𝐹 grid. This process was repeated for each frame in the event and the resulting trajectory was smoothed using a Rauch–Tung–Striebel smoother (based on the concept of Kalman filtering) as implemented in the FilterPy package for Python (https://github.com/rlabbe/Kalman-and-Bayesian-Filters-in-Python).

#### Vector field plots

Vector field plots were generated by first applying an 8 × 6 grid to the imaged region. We then pooled all event trajectories for each developmental stage and overlayed these on the grid. For each element in the grid we considered the set of trajectory segments which traversed its bounded area. We computed the velocity vector for each of these segments individually, then averaged over all segments to estimate a velocity vector for that element (i.e., the average speed and direction of propagating spatio-temporal activity in the corresponding local patch of cortex). Note that we used a grid with larger ROIs for this component of the analysis to increase the number of samples (wave trajectory segments) that were averaged to produce each vector.

#### Spatio-temporal event properties and statistics

Event distance was defined as the integral along the length of an event’s trajectory. The event size was defined as the total count of binary activations over the complete duration of an event. The event area was defined as the number of distinct ROIs for which there was at least one binary activation during the event (i.e., the total area spanned by the event as a count of ROIs).

The average direction of an event was computed as the circular mean over the set of angles along the event’s trajectory. Directional statistical tests for significant deviation from data uniformly distributed on a circle (Rayleigh, Rayleigh-bimodal, Rao spacing and Omnibus) were performed using a PyCircStat (https://github.com/circstat/pycircstat) – a Python implementation of the CircStat package (64). We implemented the Hermans-Rasson test based on (65).

#### Characterizing regional differences in neural activity

Feature difference profiles were computed for paired SS/VIS recordings. The input data were 9-dimensional feature vectors of the neural activity measures used in this study. Each feature was preprocessed by normalizing to zero mean and unit variance. Profiles were generated by subtracting each VIS feature value from the corresponding SS feature value for each animal individually, and then averaged over animals. Principal components analysis was performed on the same data using Scikit-learn. The components were evaluated on the pooled set of feature vectors from all recordings over SS and VIS.

To test for statistical differences between SS and VIS activity features at later stages, we first performed dimensionality reduction using LASSO regression to reduce the likelihood of overfitting the statistical model. To estimate the subset of features which consistently discriminated between the SS and VIS data, we regressed the 9-dimensional feature vectors against cortical region. To estimate feature importance, the model was fit for a range of the regularization parameter α, and over 100 × 5-fold repeated cross validation. We elected to keep only features with non-zero median weight in the fitted models at the optimized value of α. We did a MANOVA test on the resulting 6-dimensional feature set and visualized the separation of groups by plotting the projection of scores from linear discriminant analysis.

#### Software and code

Quantitative analysis was performed using custom code written in Python and using NumPy, SciPy, Scikit-learn, Pandas, NetworkX, Python-Louvain, Filterpy and Scikit-image. Statistical testing and modelling were performed using SciPy, Statsmodels, Pycircstat, skggm and Scikit-learn. Figures were rendered using Matplotlib. Code and associated data is available at UQ’s repository (doi.org/xxx/uql.xxx and *SI Appendix*).

### Figure preparation

Trace plot diagrams, histograms, and raster plots were all generated in Python. Image stills and standard deviation intensity projections obtained via two-photon microscopy, as well as confocal histological sections, were created and enhanced for brightness and contrast using Image J (National Institutes of Health, MD). Images obtained via fluorescence microscopy were pseudocolored, cropped, and brightness-contrast enhanced using Adobe Photoshop (Adobe Systems Inc., CA). Figures were arranged for presentation using Adobe Illustrator (Adobe Systems Inc., CA).

## Acknowledgments

We thank D. Adams, A. Deschamps, D. Herne, and C. Lau for animal support, Z. Pujic and R. Amor for assistance with two-photon imaging, and T. Holy, A. Burkhalter and D. Van Essen for critical comments on the manuscript. Live imaging was performed at the Queensland Brain Institute’s Advanced Microscopy Facility using LSM 710, generously supported by the Australian Government (LE130100078). Funding for this project was provided by National Health and Medical Research Council: GNT1159778 (LJR, GJG, RS), GNT1120615 (LJR), GNT1196855 (GJG) and 2013349 (RS); Australian Research Council: DP160103958 (LJR, RS), DE160101394 (RS) and DP200103090 (RS, LRF); The University of Queensland: Early Career Researcher Grant UQECR1719425 (RS, LA) and Postgraduate Scholarship (TB, AP); Brain and Behavior Research Foundation 26728 (RS); and National Institute of Health DP1NS127279 (LJR).

