## Supplementary data and figures for "Cortical activity emerges in region-specific patterns during early brain development"

**This PDF file includes:**

- Supporting Text
- Figures S1 to S7
- Table S1
- Legends for Movies S1 to S5

**Other Supplementary Information for this manuscript include the following:**

- Movies S1 to S5

**Supporting Information Text**

**Extended Methods**

**Animals**. All breeding and experimental procedures were approved by The University of Queensland Animal Ethics Committee and the Queensland Government Department of Environmental and Heritage Protection and were performed according to the current Australian Code for the Care and Use of Animals for Scientific Purposes (NHMRC, 8th edition, 2013). Fat-tailed dunnarts were bred at the Hidden Vale Wildlife Centre and Herston Medical Research Centre, The University of Queensland. To detect the presence of joeys in the colony, the pouches of females were inspected every two-three days. The developmental stage of joeys was therefore P0-P3 days at inspection. The developmental stage was then confirmed according to head and body milestone features.

**Plasmids.** The calcium indicator GCaMP6s (Addgene 40753, MA) was cloned into a PiggyBac plasmid (kind gift of Joseph LoTurco, University of Connecticut, USA). In this plasmid, GCaMP6s is flanked by PiggyBac inverted terminal repeats. This pPBCag-GCaMP6s plasmid was co-transfected into developing dunnarts with a pCAG-PBase helper plasmid (kind gift of Joseph LoTurco, University of Connecticut, USA).

**In pouch electroporation.** When joeys were P10-P15 (Fig. S1a), their mother was temporarily sedated by placement in a gas anesthesia induction chamber, where 5% isoflurane in oxygen was delivered at a flow rate of 200 mL/min. Sedated animals were transferred to a heating pad, and anesthesia was maintained via 5% isoflurane in oxygen delivered through a silicone mask (Zero Dead Space MINI Qube Anaesthetic System, AAS, AZ). The joeys were exposed without removing them from the teat by carefully everting the pouch, and 0.5-1 μL of a solution containing 1 μg/μL of pBase plasmid and 1.5 μg/μL of GCaMP6s plasmid in 1M phosphate buffered saline (PBS; Lonza, Basel) and 0.0025% of the dye Fast Green (Sigma-Aldrich Co., MO; to visualize the location of injection) was injected into the right lateral ventricle using a pulled glass pipette (WPI, FL) attached to a picospritzer (Parker Hannifin, NH). Five square pulses of 100 ms duration and 35 V were delivered at 1 Hz using 3 mm tweezer-like electrodes (Nepa Gene Co., Ichikawa) via an electroporation system (ECM830, BTX, Harvard Bioscience, MA), with the positive electrode placed above the developing cortex. In order to identify electroporated animals at later stages, animals were tattooed by scratching their paws with a hypodermic needle filled with green tattoo paste (Ketchum Mfg. Co., NY).

**Surgical procedure for *in vivo* imaging.** Joeys were imaged between P20-P40. Mothers were anaesthetized and joeys were exposed as described above. Joeys were carefully removed from the teat by gently pulling, whilst squeezing the base of the teat with forceps. The joey was then restrained by wrapping its body in gauze and encasing it within a modified 1.5 μL centrifuge tube (Eppendorf, Hamburg), leaving the head exposed. Joeys were then anesthetized on ice for 4-6 minutes, or until unresponsive to stimuli. Topical local anesthesia (1% lignocaine; AstraZeneca, Cambridge) was applied to the skin on the scalp with a fine paintbrush, and the scalp covering the electroporated area was then removed. It was also critical to carefully remove the muscles superficial to the skull to prevent contractions that moved the sample during calcium imaging. Finally, a glass-bottomed petri dish (MatTek, MA) was fixed to the skull of the dunnart joey with cyanoacrylate glue (UHU, Bühl). The animal was then allowed to recover from ice anesthesia whilst the cyanoacrylate set for at least 30 mins.

**Figure preparation**. Trace plot diagrams, histograms, profile and raster plots were all generated in Python. Image stills and standard deviation intensity projections obtained via two-photon microscopy were created and enhanced for brightness and contrast using Image J (National Institutes of Health, MD). Images obtained via fluorescence microscopy were pseudocolored, cropped, and brightness-contrast enhanced using Adobe Photoshop (Adobe Systems Inc., CA). Figures were arranged for presentation using Adobe Illustrator (Adobe Systems Inc., CA).

**Figures S1 to S7**

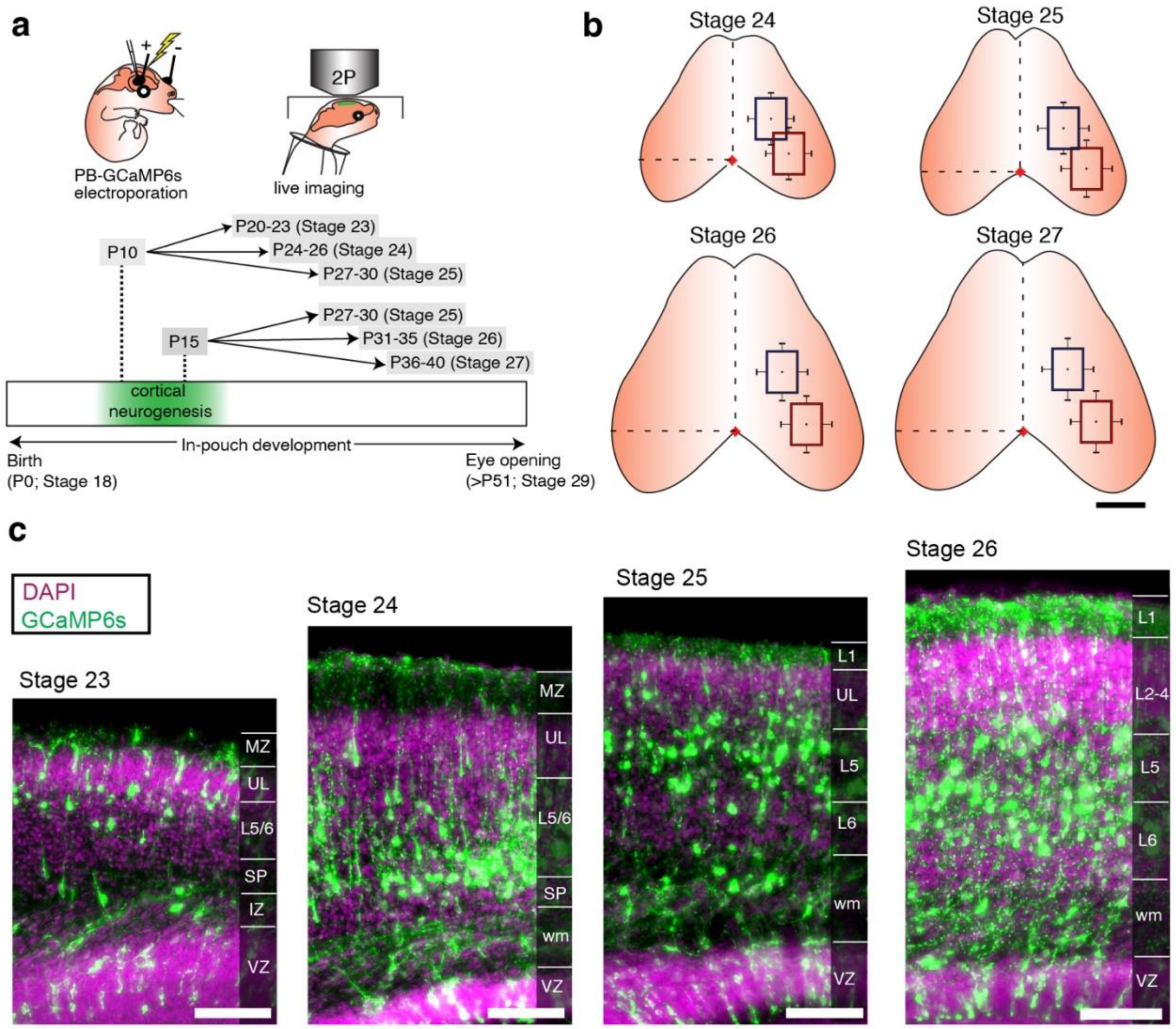


**Figure S1: Transfection, live imaging and histology of labelled neurons across stages.**

**a,** In-pouch electroporation of PB-GCaMP6s was performed during the peak of cortical neurogenesis, at postnatal days (P) P10 or P15 for subsequent live 2P-imaging at stages 23-25 and 25-27, respectively. **b,** The average position of the imaged areas and standard error is indicated in blue and red for SS and VIS, respectively, at the indicated stages, relative to lambda (red dot). See Table S1 for values. **c,** Distribution of GCaMP6s expression (green) in cortical neurons across stages 23 to 26. Note distribution of GCaMP6s cell bodies across layers (L) and dendritic arborizations within the marginal zone and layer 1. IZ, intermediate zone; MZ, marginal zone; UL, upper layers; VZ, ventricular zone. Scale bar 1 mm in **b,** 50 µm in **c**.

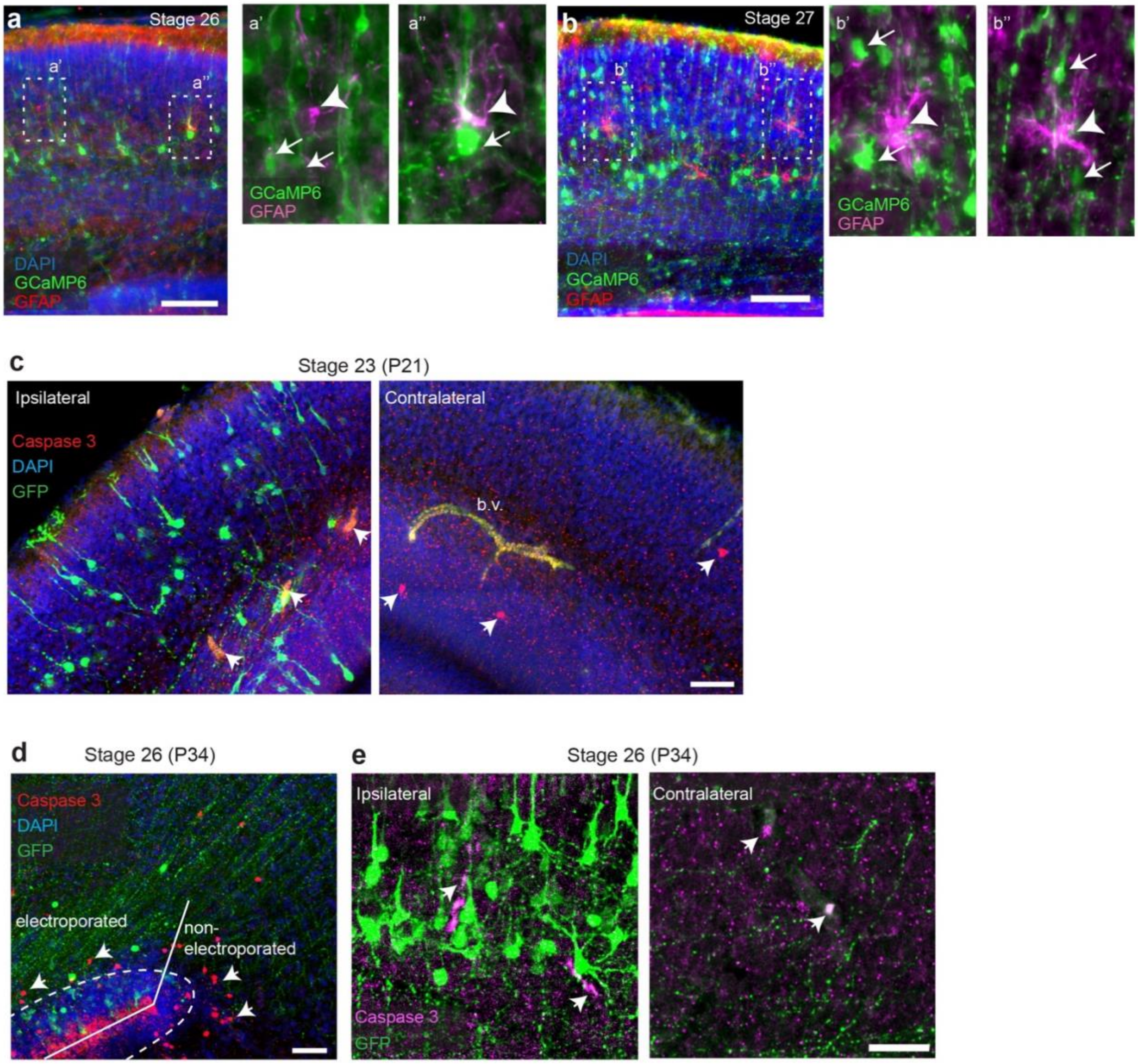


**Figure S2: GCaMP6s cells do not express glial marker GFAP nor induce cell death.**
**a**, GFAP-positive cells (red) are very few and sparse in the developing cortex by stage 26. Insets a' and a'' show higher magnification of the dotted regions on left, with a few GFAP-expressing astroglia (arrowheads, magenta) not colocalizing with GCaMP6s neurons (arrows, green). **b**, Glial cells become more prominent from stage 27 onwards, and do not colocalize with GCaMP6s, as seen in insets on right. **c-e,** The cell-death marker cleaved caspase 3 was very sparsely expressed from stage 23, in similar numbers between electroporated and contralateral hemispheres (arrowheads). **d-e,** Similar extents of caspase 3 expression (red) within electroporated and adjacent non-electroporated basal areas (**d**), and within upper-layer electroporated cells and contralateral regions (**e**), suggesting that cell death is not a major influence in our experimental paradigms. b.v., blood vessels. Scale bars: 50 µm.

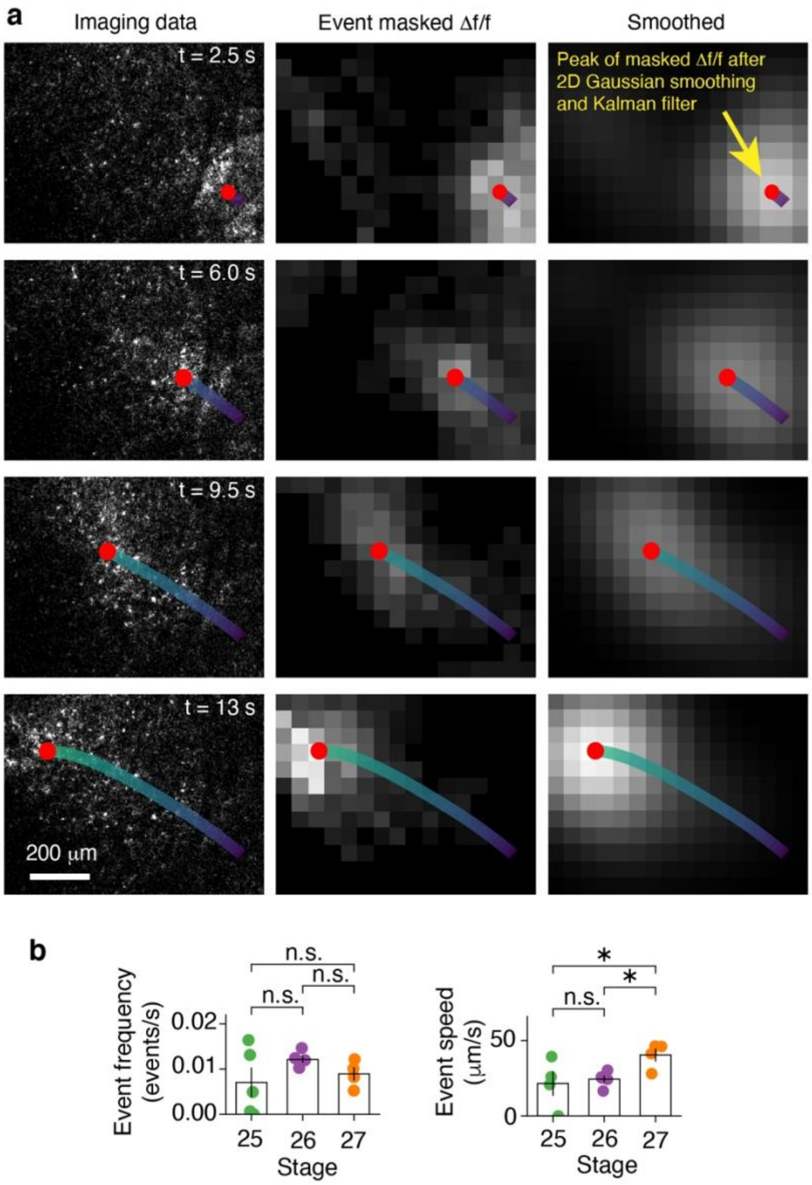


**Figure S3: Automated detection and tracking of waves.**

**a**, Illustration of the wave tracking algorithm (see Extended Methods). The 2-dimensional spatial array of ΔF/F time series for a given frame was masked to retain contiguous blocks of activity (significant deviations above baseline fluorescence) which were greater in size and duration than would be expected by chance. For each distinct event, the resulting event-masked image was used to identify the spatial peak in activity for each frame. The trajectory of this point through time was then smoothed using a Kalman filter. **b**, Detected event frequency does not change significantly over development. While seemingly at odds with the overall increase in activity over development, this result is likely explained by the method used to set thresholds for event detection – a percentile applied to connected component statistics on temporally shuffled data – which will typically produce a higher threshold for above-chance events when there are more significant deviations above baseline fluorescence, regardless of whether the increase in such deviations was due to noise or genuine neural activity. Event speed increases significantly between stage 26 and 27.

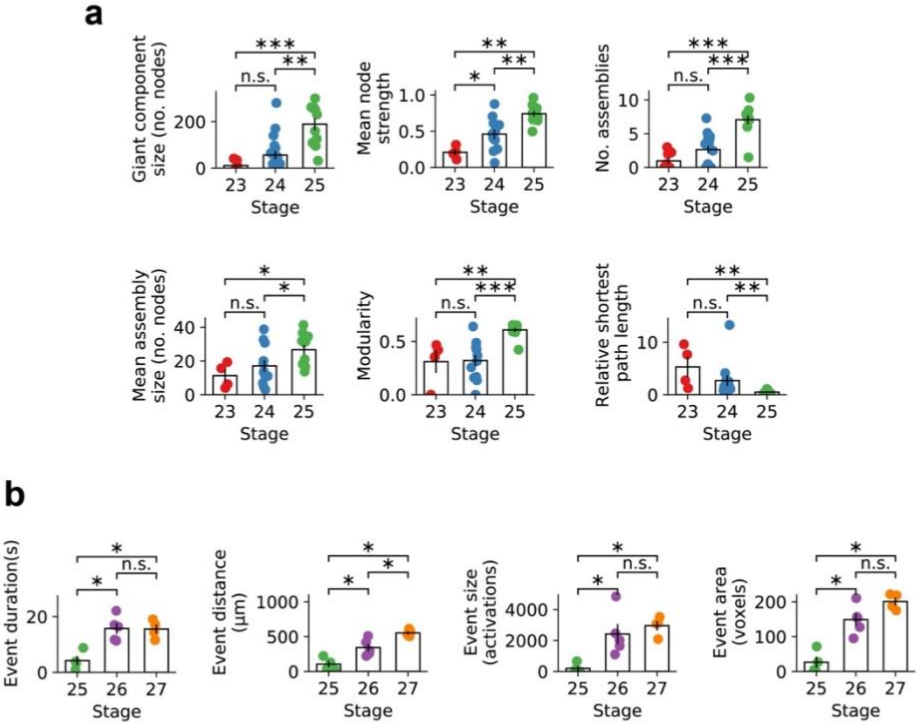


**Figure S4: Reduced size ROI analyses does not alter the main findings in SS or VIS**.
The onset and maturation of patchwork and wave activity in the dunnart somatosensory (**a**) and visual (**b**) cortices. These analyses were performed in the same way as those in Figs. 2 and 3, but using regions-of-interest (ROIs) with size linearly reduced by 20% (e.g., approximately 2/3 of area). All trends and statistical results are consistent with the results reported in the main text, with slight differences in p-values.

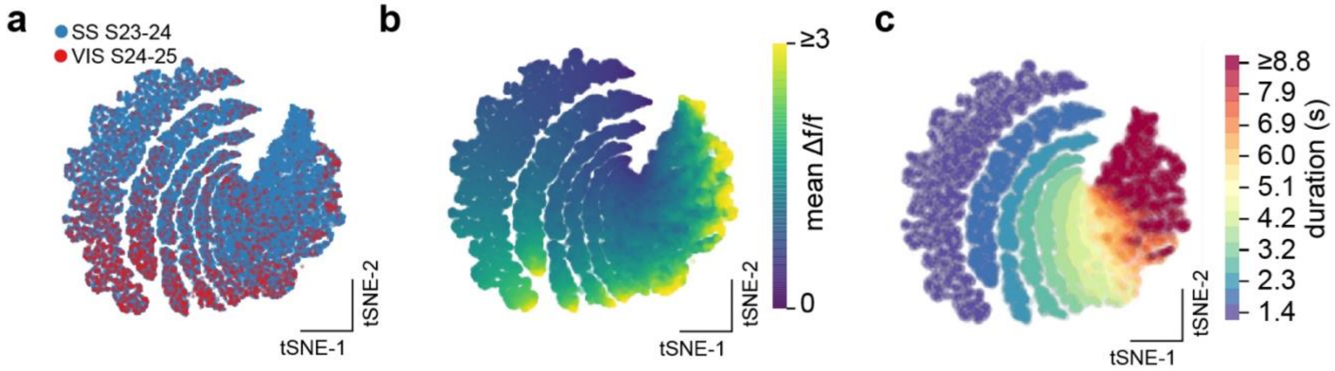


**Figure S5: Local calcium transients differ between regions from earliest pre-onset and onset activity**. **a-c**, t-SNE embedding of individual calcium transients from single ROIs in somatosensory (SS) and visual cortices (VIS) for recordings from Stages (S) 23-25, corresponding to pre-onset and onset sparse activity (see Fig. 4c). Each point represents an individual calcium transient. Data were first padded to a fixed length, then principal component (PC) analysis was applied to reduce dimensionality to 10 PCs explaining ≈93% of the variance in the data. Individual events segregate in the low-dimensional representation, as shown in (**a**) color-coded by region (SS, blue, VIS, red), suggesting differences in the nature of neural activity at a local scale. **b,** All data color-coded by the mean Δf/f of each event (approximation of the area under the curve relative to the duration) also indicates differences, which are quantified by area as shown in Fig. 4d. **c,** Same data as in (**a**) with each event color-coded by duration in seconds (s) reveals discrete stripes of increasing duration from left to right. This striped structure is very-likely to be a non-biological artefact resulting from discrete-time sampling of the data. We have only considered events that span at least 3 time points (≈1.4 seconds).

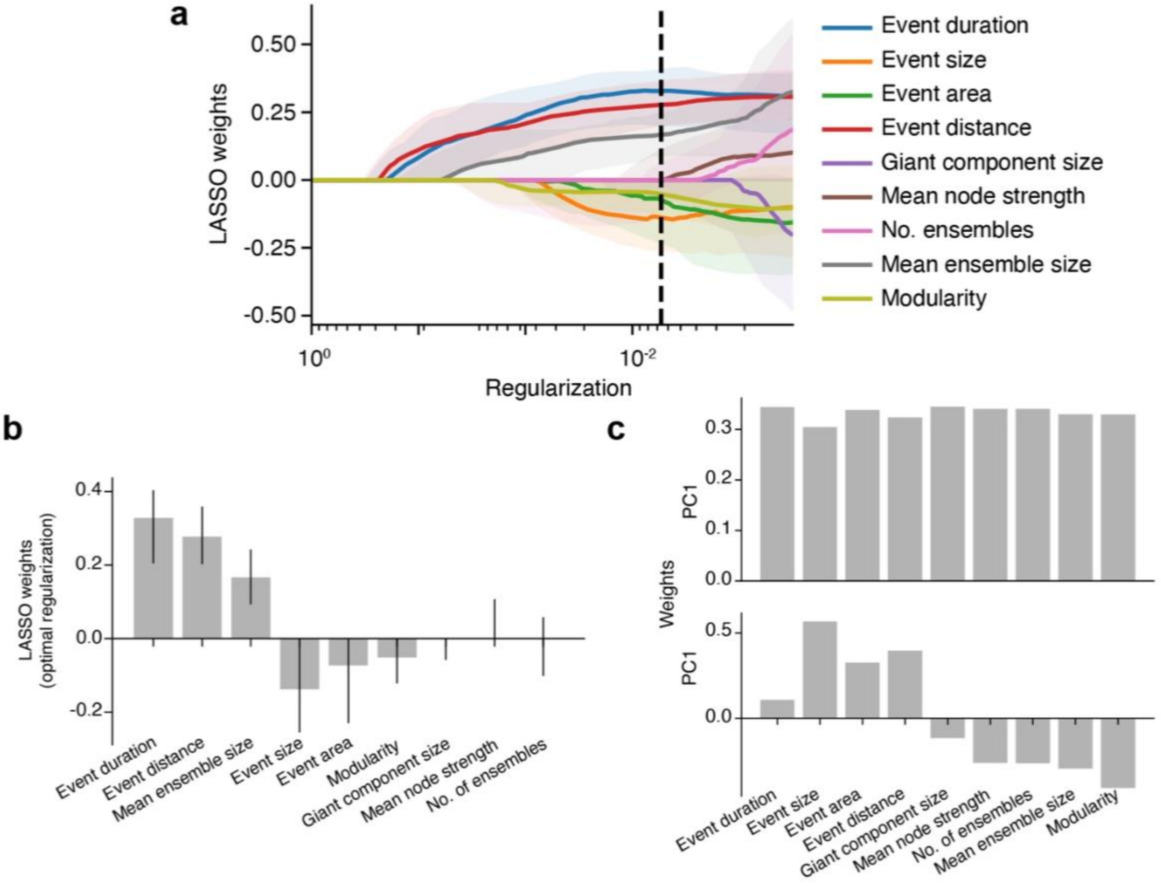


**Figure S6: Principal component analysis of neural activity features and dimensionality reduction using LASSO regression.**

**a,** The regularization path for LASSO regression on neural activity features from pooled data at stage 26 and stage 27 from SS and VIS. Colored curves and shaded regions show median feature weights and 95% confidence intervals respectively over 100 repetitions of 5-fold cross validation. The regularization parameter controls model complexity to minimize model error while reducing the likelihood of overfitting. In LASSO regression, decreasing the regularization parameter sequentially switches on features in the model one at a time in the order that best improves model performance. The vertical dashed black line indicates the optimal regularization parameter for the data. We reduced the size of our feature set for subsequent statistical testing by only including features with non-zero median weight for the optimal regularization. **b**, Median weighting and 95% confidence interval for each feature using from LASSO regression for the optimal value of the regularization parameter (corresponding to the dashed black line in **a**). **c**, Feature weights for the first two PCs between areas and across stages 24-27, as plotted in Fig. 5b. Top plot represents increasing values for all features, which characterize patchwork and wave activity respectively. The negative direction of the average feature difference between SS and VIS for early-stage recordings is congruent with the delayed onset of activity in VIS.

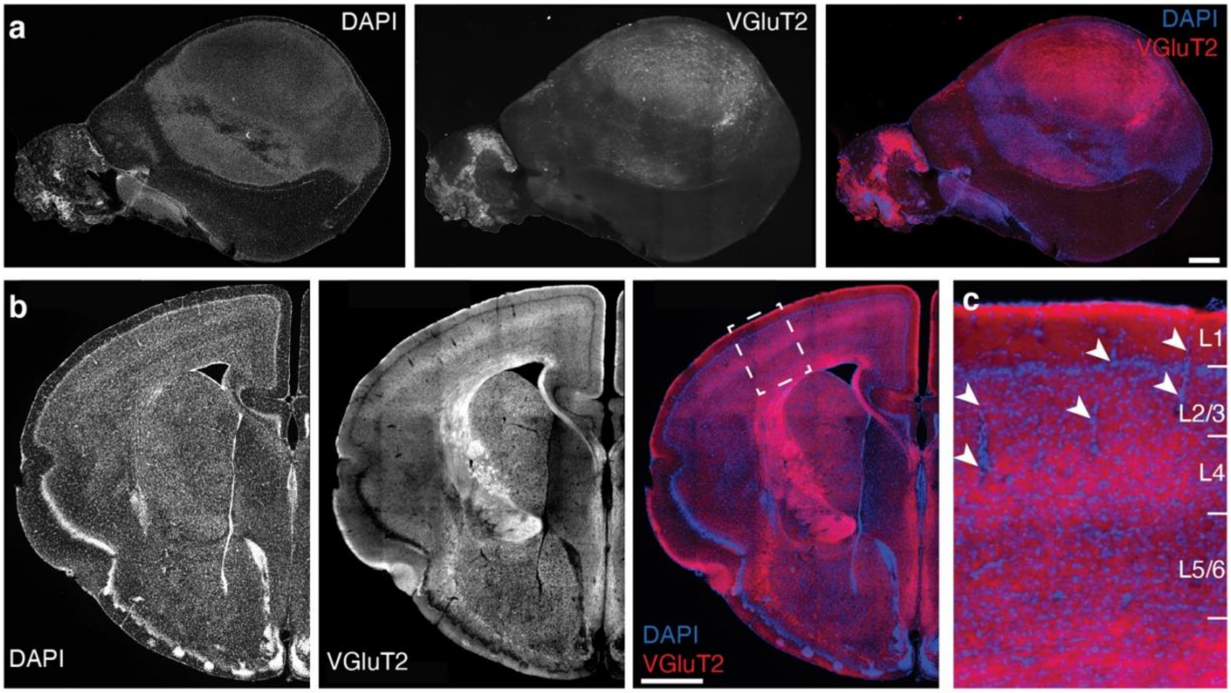


**Figure S7: Lack of histologically evident barrels in SS of adult dunnarts.**

**a**, Tangential flat-mount section of an adult dunnart brain at the level of layer (L) 4, as indicated by DAPI staining (left) and immunofluorescence against the thalamocortical axon marker VGluT2 (middle). Right panel is a merge of both channels revealing thalamocortical innervation with the absence of cortical barrels. **b**, Coronal brain section of an adult dunnart across SS showing a VGluT2-dense layer 4 that lacks barrels and extends into adjoining cortical areas. The square region in the merged panel is expanded in **c** revealing the continuity of L4, and some blood vessels (arrowheads) across layers. Scale bars: 1000 µm in **a** and 500 µm in **b**.

**Table S1.**

Summary of the coordinates of the center of imaged areas, relative to lambda, of somatosensory (SS) and visual (VIS) areas across developmental stages (S). Data in µm ± S.E.M.

| | S24 | | S25 | | S26 | | S27 | |
|---|---|---|---|---|---|---|---|---|
| | Rostral | Lateral | Rostral | Lateral | Rostral | Lateral | Rostral | Lateral |
| SS | 850 ± 100 | 800 ± 150 | 900 ± 100 | 900 ± 200 | 1200 ± 150 | 900 ± 200 | 1250 ± 200 | 900 ± 200 |
| VIS | 150 ± 100 | 1150 ± 150 | 100 ± 150 | 1300 ± 200 | 150 ± 150 | 1450 ± 200 | 200 ± 200 | 1500 ± 200 |

**Movie S1 (separate file).**

Patchwork neural activity imaged from SS of dunnarts at stage 26 (postnatal day 32).

**Movie S2 (separate file).**

Travelling wave activity imaged from VIS of dunnarts at stage 26 (postnatal day 32).

**Movie S3 (separate file).**

Automated tracking of wave trajectories in VIS from a dunnart at stage 27 (postnatal day 39).

**Movie S4 (separate file).**

Onset of patterned neural activity imaged from SS of dunnarts at stage 24 (postnatal day 24).

**Movie S5 (separate file).**

Onset of patterned neural activity imaged from VIS of dunnarts at stage 25 (postnatal day 28).
